# Hypothesis-driven modeling of the human lung-ventilator system: A characterization tool for Acute Respiratory Distress Syndrome research

**DOI:** 10.1101/2022.10.31.514563

**Authors:** J.N. Stroh, Bradford J. Smith, Peter D. Sottile, George Hripcsak, David J. Albers

**Author notes:** Corresponding author *Email address* (J.N. Stroh).

## Abstract

Mechanical ventilation is an essential tool in the management of Acute Respiratory Distress Syndrome (ARDS), but it exposes patients to the risk of ventilator-induced lung injury (VILI). The human lung-ventilator system (LVS) involves the interaction of complex anatomy with a mechanical apparatus, which limits the achievable flexibility and fidelity needed to provide individualized clinical support by modeling lung processes. This work proposes a hypothesis-driven strategy for LVS modeling, in which robust personalization is achieved using a pre-defined parameter basis in a non-physiological model. Model inversion, here via windowed data assimilation, forges observed waveforms into interpretable parameter values that characterize the data rather than quantifying physiological processes. Inference experiments performed on human pressure waveform data indicate the flexible model accurately estimates parameters for a variety of breath types, including breaths of markedly dyssynchronous LVSs. Parameter estimates generate static characterizations of the data that are 50–70% more accurate than breath-wise single-compartment model estimates. They also retain sufficient information to distinguish between the types of breath they represent. However, the fidelity and interpetability of model characterizations are tied to parameter definitions and model resolution. These additional factors must be considered in conjunction with the objectives of specific applications, such as identifying and tracking the development of human VILI.

## 1. Introduction

Hospitalized patients may require assisted ventilation, either due to the inability to breathe on their own or because care providers determine they should not do so. In such cases, respiration is supported by a programmable mechanical ventilator that actively supplies air to the lungs. Proper ventilator management is a particular concern in patients presenting acute respiratory distress syndrome (ARDS), a life-threatening condition associated with mortality in: 3–4% of ICU admissions [1, 2] and nearly 60% of ventilated COVID-19 patients under intensive care [3]. Mismatch in timing, breath-trigger threshold, or pressure/volume budget evince dyssynchronies between patient and machine, which may contribute to problems such as edema and hypoxia [4]. Further, ventilatory dyssynchrony can cause parenchymal damage termed ventilator-induced lung injury (VILI) and may exacerbate the effects of ARDS. Clinical management, therefore, requires carefully specified thresholds and target quantities to provide necessary and sufficient respiration[5] while avoiding VILI, a major risk for assisted patients[6]. The prevalence of these cases strongly motivate additional lung-protective considerations [7, 8, 9, 10] to improve patient outcome.

Respiratory management would benefit from an improved understanding of the lung within the context of the clinical environment where VILI may occur, which includes the ventilator component. Many investigations use controlled ventilation to probe lung properties ([11, 12, 13, 14, 15] as well as [16] and references therein). Compartment-based models, described in the following subsection, estimate the observed pressure (*p*) and volume (*V*) relationship in terms of lung compliance and resistance. However, observations of airway pressure and volume (*p* and *V*, respectively, or *pV* together) largely originate in the ventilator and are observed outside the patient to reflect the bulk effect of the lung system. Consequently, these existing models try to infer specific parameters governing lung dynamics from observations of aggregate respiratory, systemic, and healthcare processes.

The lung-ventilator system (LVS, comprising the interaction of the lungs and breathing support) exhibits a variety of waveform dynamics that are of primary concern in the study of VILI and ventilator dyssynchrony [17, 18, 19]. Capturing these behaviors in process-oriented models requires a high degree of either complexity or parameterization, with additional overhead needed to account for heterogeneity among patients and ventilation therapies. Methods for identifying, classifying, and preventing dyssynchronous timing and delivery of breath support are nascent applications in clinical informatics.

This work focuses on a hybrid model-based method for transforming LVS data into discrete parametric summaries that retain features of interest. The intended use is in informatics applications requiring description and categorization of the LVS, rather than material lung properties, for further analysis in the context of health care process information such as ventilator settings. Such a tool would benefit the scientific inquiry of VILI by facilitating data representation in algorithms capable of detangling the effects of ventilator settings from changes in the lung response. The following subsection revisits the existing method of LVS estimation to motivate the proposed approach oriented toward operational LVS representation and clinical translation.

### 1.1. The dynamic single-compartment model: framework and limitations

The single-compartment model ([20]) defines a stationary relationship of *pV* in the respiratory tract by the equation

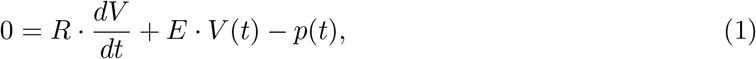

with parameters (possibly nonlinear functions) for resistance (*R*) and elastance (*E*). In practice, *p* is the pressure with respect to a zero-volume reference; the system requires a baseline pressure parameter (viz. positive end-expiratory pressure, or PEEP) for the relationship to hold for arbitrary initial times.

This model is a fast and efficient means of estimating physiological lung properties from the *pV* relationship. However, requires higher-fidelity process resolution to capture features of clinical interest even for the simplest cases [21]. Fig1) illustrates the failure of the linear model to capture waveform features such as the pressure plateau shape related to the alveolar recruitment rate. Nonlinear extensions widen resolvable features by adopting time- or state-dependent sub-models of tissue rheology to model specific components of lung injury from mechanistic understanding [21, 22, 23, 24, 25, 26, 16], while others increase complexity using multi-compartment frameworks [27, 28, 29, 30]. These augmentations resolve additional processes but impose additional limitations, and no identified literature reports their application to mechanically-ventilated human data.

**Figure 1:**
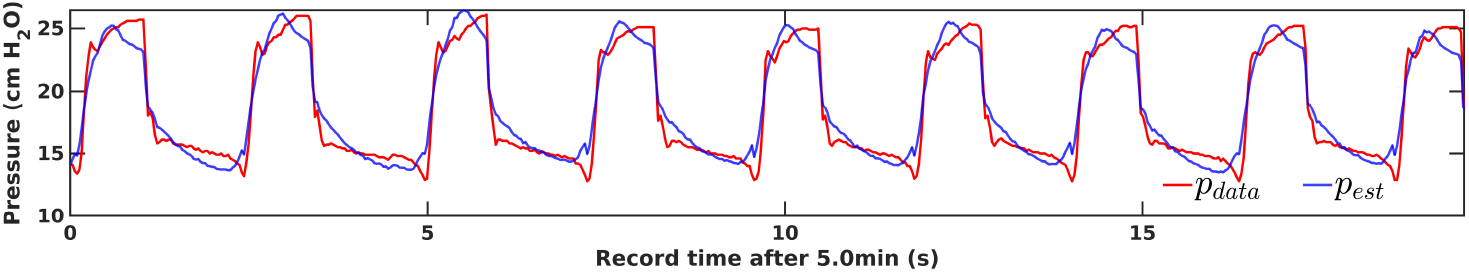
Linear single-compartment state resolution: pressure observation (*p*_data_) vs. optimal single compartment estimation (*p*_est_) for human breaths of patient #1 with mild ventilator dyssynchony [19]. Information about lung damage and disease may manifest itself in the shapes of waveform features not resolvable by the assumed *pV* relationship.

Limits of the compartmental framework create obstacles to personalization and clinical translation. The increased parametrization required to resolve specific dynamics may not capture signatures of ventilator dyssynchrony, such as those discussed in [31, 32, 33], in a way easy to estimate from data. Nonlinear compartmental models, such as those with state-dependent parametric effects, are more difficult to invert due to strongly correlated state and parameter effects. Inversion of multi-compartment models may not uniquely identify and ascribe parameter values without additional LVS knowledge even when compartments have unique parametric timescales. Likewise, time-varying formulations [34, 35, 24] may fully resolve dynamic elastance without an assumed structure needed to convey this information compactly. Caregivers versed in waveform descriptors and ventilation annotation schemes [36] may also have difficulty adopting complex parametric descriptions of lung deformation properties. Such ontological differences further limit the scope and effectiveness of translating patient-specific parameters into clinical and translational informatics (e.g., machine learning) domains for desirable applications such as LVS phenotyping and classifying dyssynchronous behaviors [19].

### 1.2. Purpose & Outline

Clinical informatics is the appropriate domain for identifying and understanding the incidence of VILI, as it involves quantifying the effects of the healthcare process. However, this first requires a flexible and compact description of LVS features from heterogeneous data that existing models do not provide. This work presents an inferential modeling approach to robustly describe patient-specific LVS systems with interpretable parametric descriptors, where expert knowledge or other a priori definitions replace theory-driven mechanistic processes in the dynamical system. At the inference level, proposed parameter values correspond to hypotheses that are refined to fit observed data. Higher-level scientific hypotheses regarding VILI and LVS systems may then be pursued, leveraging these parametric representations of data in downstream informatics processes. The central hypothesis in this work is that parameter-expressed waveforms retain interpretable, distinguishing characteristics of data in digitized form, properties essential to its use in such applications.

The remainder of this work is outlined as follows. Section 2 proposes a model approach to parametric estimation of LVS waveform data, outlines the employed inference scheme, and describes the data source. Section 3 presents experimental results of parameter estimation on human *pV* collected from ventilated patients diagnosed with ARDS. Section 4 discusses the gains and limitations of the proposed methodology in application to the clinical estimation environment.

## 2. Method

This section presents an LVS estimation approach comprising a simple linear dynamical system (Sec. 2.1) combined with a windowed ensemble-based estimation scheme (Sec. 2.2). The dynamical model generates waveforms from a simple, parametrically-defined function that can be data-optimized to yield a parametric form of the ingested data. A windowed ensemble Kalman- type smoothing method is employed here to assimilate patient data and systematize *pV* data into a small set of patient-specific descriptors.

### 2.1. The data representation model

The presented model determines a parametric representation of LVS waveforms using a method conceptually recent related to work recent [37]. Rather than simulating underlying data-generating processes, it extracts interpretable descriptors from LVS data applicable to a range of behaviors including pathophysiological cases such as dyssynchronous breaths. The approach circumvents direct resolution of the lung and ventilator which may require many parameters to model robustly, involve bulk process parametrizations at some scale, and must ultimately sacrifice mechanistic fidelity for functionality.

The proposed LVS waveform model simulates a signal *x*(*t*) (e.g., pressure or volume) according to the equation:

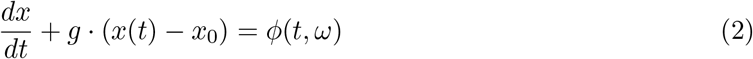

where *g* > 0 is a smoothing parameter and *ϕ* is a time-dependent function of parameters *ω*. The value *x*_0_ corresponds to a reference baseline, such as ventilator PEEP in LVS pressure applications. The user-defined periodic function *ϕ* drives the system dynamics and is referred to as the forcing. The model transforms parameters defining the function *ϕ* into the continuous state output *x*. The forcing and output states have equivalent parameter dependencies within this linear differential equation, with the parameters giving rise to localized features of the solution. In the following applications, the respiratory cycle length *θ* and baseline value *x*_0_ are fixed values but may be estimated in other applications targeting patient-triggered ventilation.

The relationship of parameters to the solution is specified by forcing function *ϕ*, which may be application-specific. One may target specific waveform features or particular nuances of interest for a given application, leveraging domain-expert knowledge to maximize the relevance of parametric representation. For simplicity, presented applications use a step function *ϕ_M_* obtained by discretizing the breath cycle into *M* equal epochs:

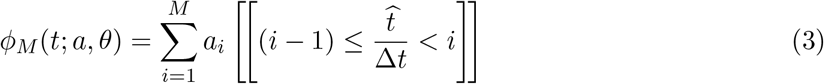

where *θ* is the breath cycle duration, 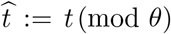 is local breath time, *i* = 1..*M* indexes the epochs with common length Δ*t* ≔ *M/θ*, *a* is a vector of forcing amplitudes, and ⟦·⟧ is the logical operator. The number of epochs-per-breath *M* defines the model resolution, and the simplified model is identified by the symbol *ϕ_M_* in this discussion.

Interpretability is inherited through a priori parameter definitions because the parameters as defined by Eqn(3) have non-overlapping domains. Parameter values are therefore directly associated with the waveform amplitude and derivative over the time epochs that define them. In the LVS case, the first parameter is associated with initial inspiration and the last with the end of a breath; the other parameters have a resolution-dependent association with ordered epochs elapsing during the breath sequence. In the generalization of Eqn(3) using unequal epochs, redefining the amplitudes as

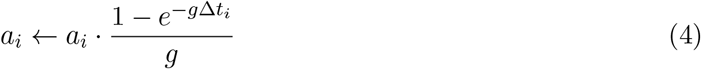

balances the estimability of the parameters by making their magnitudes independent of epoch lengths.

### 2.2. Parameter Inference via data assimilation

Personalizing the model proceeds by optimizing parameters so that simulated output fits the target data. Tracking the LVS evolution of a patient under mechanical ventilation, therefore, requires estimating *O*(10^5^) breaths per day. The need to estimate short-time features over this much longer scale constrains inference options based on speed, robustness, and comparability of estimates; such considerations discourage the use of Markov Chain Monte Carlo methods and windowed spectral decomposition.

In this work, parameter inference is accomplished by a windowed ensemble Kalman smoother (EnKS) [38, 39]. It approximates a Bayesian update scheme by identifying parameter optima using the error structure of perturbed ensemble simulations over a window (*t*_0_, ..*t_k_, …t_L_*] to minimize difference between the ensemble-mean forecast and associated external data 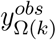 (*k* = 1..*L*). Formally, this is achieved by identifying the augmented model state (state + parameters) that minimize the cost function

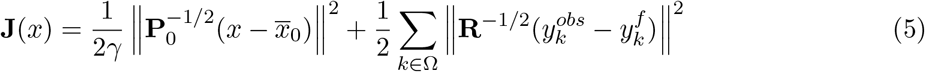

where *γ* is a covariance inflation factor to control the weight of data; initial data for the ensemble are characterized by mean 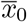 and covariance 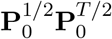; and **R**^1/2^ is the observation error covariance matrix. Symbols 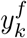 represent simulated observations based on initial state *x*, and Ω is a sub-indexing of time to regulate assimilated observations. Parameter variations about the optimal mean are retained to re-initialize the forecast at a point *t_S_* within the forecast window. Fig2 illustrates the inference scheme, with additional details in SI. Appendix A.

**Figure 2:**
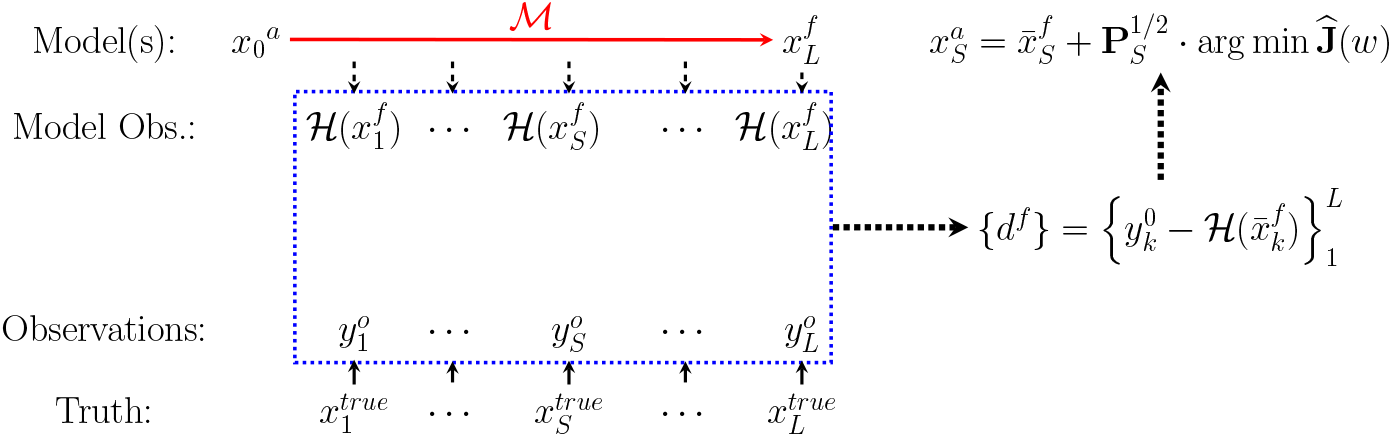
The schematic windowed assimilation process. The conceptual relationship between the true state, observational data, modeled data, and model states is shown. The model (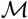, red) performs the forecast process over the window (blue), simulating equivalents of data as well as the error covariance structure. The optimal solution at any time *t_S_* within the window derives from error analysis of data-forecast mismatch.

### 2.3. Experiments and Data

The following section reports the inference of model parameters in experiments applied to both synthetic and clinical human data. Artificial waveform data are generated by the model using prescribed parameter values. These experiments verify the method against a known solution with nonstationary parameters that affect the solution waveform shape. Further experiments use continuous 40-breath sequences from human subjects diagnosed with ARDS under volume-controlled ventilation. These sequences contain identifiable breath types that lack artifacts (i.e.. discontinuities) related to ventilator data output and have relatively uniform waveforms within them. Each sequence is composed of either normal breaths or dyssynchronous breaths identifiable as flow-limited (FL), early ventilator-termination (eVT), or early reverse-trigger (eRT) [32]; see [19] regarding the mechanics and clinical implications of these dyssynchronies.

## 3. Results: Inferential data parametrization

This section explores experiments inferring parameter representations of data using the proposed inferential modeling method presented above.Data are estimated by optimizing model parameters, and these estimates are statistically summarized over each 40-breath sequence of data. Characterization of the data, a static waveform representative, is generated by applying the forward model to the summary mean (or another estimator). Supplemental Appendix C reports on synthetic data experiments conducted to verify the model inversion against solutions with known parameters. Those results indicate that estimated parameters quickly and accurately respond to changes in assimilated data, indicating its suitable application to the LVS system. The ensemble approximation also captures the parameter covariance structure (*viz*. the upstream dependence of *a_i_* on *a_i_*_−1_) that may otherwise confound inversion. The following sections estimate human LVS data in normal and more pathological cases, including parametric representation of pressure-volume loops. Supplemental Table B.3 contains experiment level hyper-parameters.

### 3.1. Normal breath estimates

In this and the following section, experiments explore 40-breath sequences of clinical LVS data from patients with ARDS. These 80–100 second intervals illustrate the fidelity of estimates for breaths of similar type, although practical applications may require shorter intervals for less regular data. Table 1 presents breath-level mean errors for the pressure and volume estimated by parameters at various model resolutions for the cases presented. Experiments focus on pressure waveform estimation because the ventilator controls the volume delivery throughout each experiment. PEEP values (*x*_0_) for pressure data are estimated as the mean minimum pressure across all breaths, while volume applications assume a zero-valued baseline. Period regularization within each window maintains a consistent parameter definition throughout the breath sequence and, in practice, may be applied to shorter sub-sequences when the breath cycle is irregular.

**Table 1:** Breath-averaged RMS differences of continuous pressure estimation for human experiments, with the lowest value for each sequence in bold.

| Model: | $\phi_{12}$ | $\phi_{18}$ | $\phi_{24}$ | $\phi_{30}$ |
| --- | --- | --- | --- | --- |
| Pressure (cm H<sub>2</sub>O) |  |  |  |  |
| p#1 Normal 1 | 0.24 | 0.24 | **0.24** | 0.24 |
| p#1 Normal 2 | 0.25 | 0.25 | 0.25 | **0.25** |
| p#2 FL | 0.10 | 0.10 | **0.10** | 0.10 |
| p#2 eVT | **0.13** | 0.14 | 0.14 | 0.14 |
| p#2 Normal | 0.29 | 0.29 | 0.29 | **0.29** |
| p#2 eRT | 0.34 | 0.34 | 0.34 | **0.34** |
| Volume (ml) |  |  |  |  |
| p#1 Normal 1 | **4.39** | 4.40 | 4.41 | 4.42 |
| p#1 Normal 2 | 4.09 | 4.11 | 4.13 | **4.06** |
| p#2 FL | 4.46 | **4.46** | 4.49 | 4.59 |
| p#2 eVT | 4.25 | 4.24 | **4.24** | 4.48 |
| p#2 Normal | 4.01 | 4.00 | 4.00 | **3.92** |
| p#2 eRT | **3.14** | 3.18 | 3.24 | 3.33 |

Fig3 shows the inference process using 12 epochs per breath (*ϕ*_12_) for sequences of normal, mildly dissimilar breath waveforms in patient#1 recorded 8.4 days apart. Individual breaths are well-resolved in continuous time with small relative errors that occur when the epoch-based parameter definitions span breath features. Inferred parameters give a low-dimensional representation of the data as both parameter vectors and state characterizations, together with quantified parameter uncertainties. The accuracy of these model summaries is affected by the parameter definitions, with characterized waveform including artifactual signatures of the parametric resolution.

**Figure 3:**
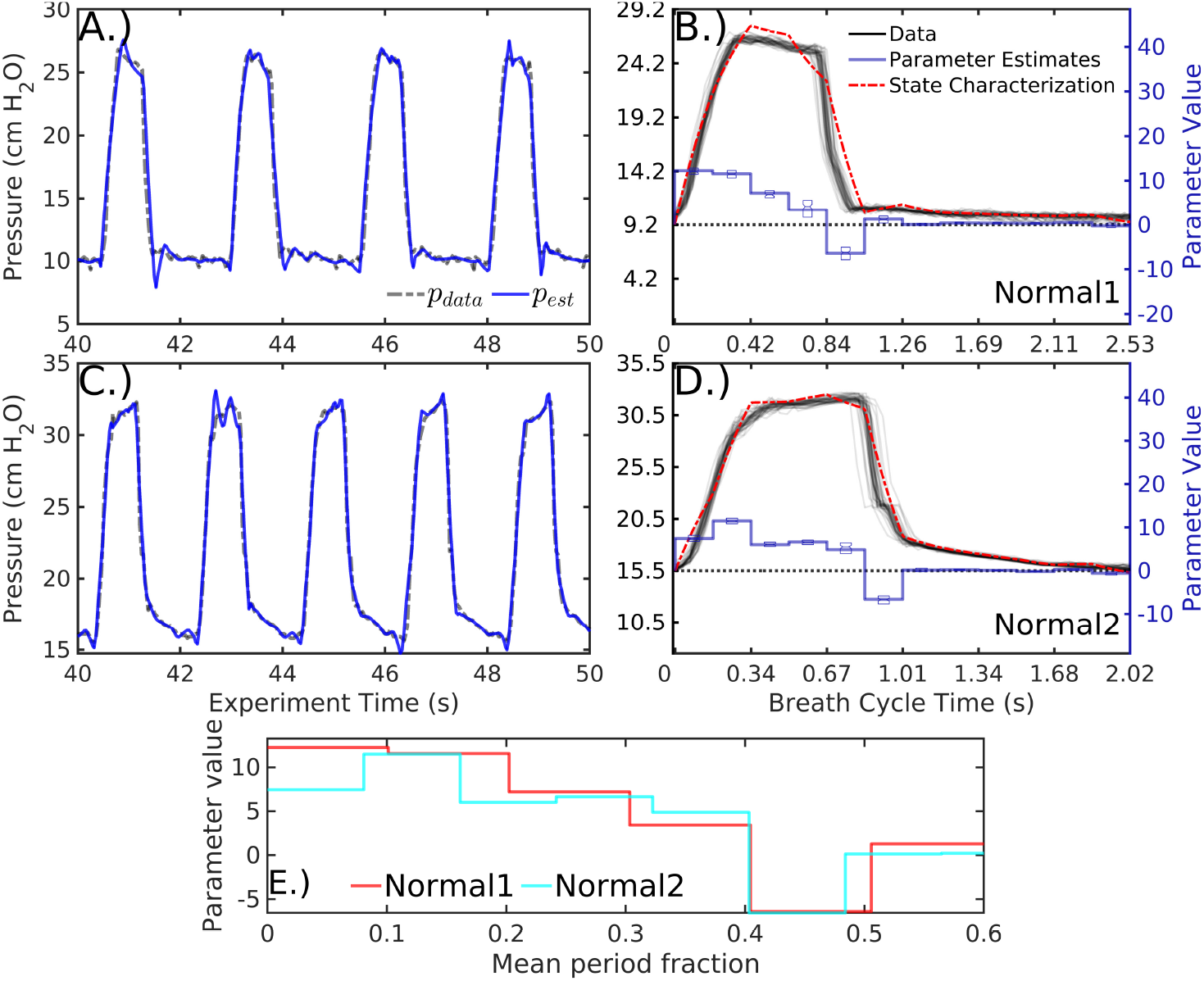
Comparing two sequence characterizations of patient#1 pressure data with 12 parameters *ϕ*_12_. Data (A and C, dashed black) are well-estimated by inference model estimates (blue) in continuous time on overlapping 1.2 second analysis windows. Tabulating data (B and D, black) and mean parameter estimates (blue, with inter-quartile range shown) in cyclic breath time compares the characterization (red) of each sequence with the data. Mean parameter vectors of the two sequences, which have significant distributional differences, are depicted over a normalized breath duration (E).

Parameter estimate distributions of the two sequences of normal breaths distinct (*p* < 0.05, approximated via [40]), providing a direct way to categorize and distinguish breath sequence types. Differences in parameter mean likewise coarsely resemble differences in the breath shape. For example, the inspiration parameters of the first sequence during the first half of the breath show a monotone decrease whereas the second sequence’s parameters rise later into the breath, just as in the waveform pressure shapes (Fig3E).

### 3.2. More complex varieties of human breaths

Patient#2 data comprise four sequences of breath types most likely identified as flow-limited (FL), early ventilator-termination (eVT), normal, and early reverse-trigger (eRT). The sequences occur 4, 3, and 14 hours apart, respectively, and their nearly equal periods permit a more direct parametric comparison. Figure 4 shows the characterization and mean summaries of these sequences as in Fig3. Parameter distributions are again distinct (*p* < 0.05 pairwise).

**Figure 4:**
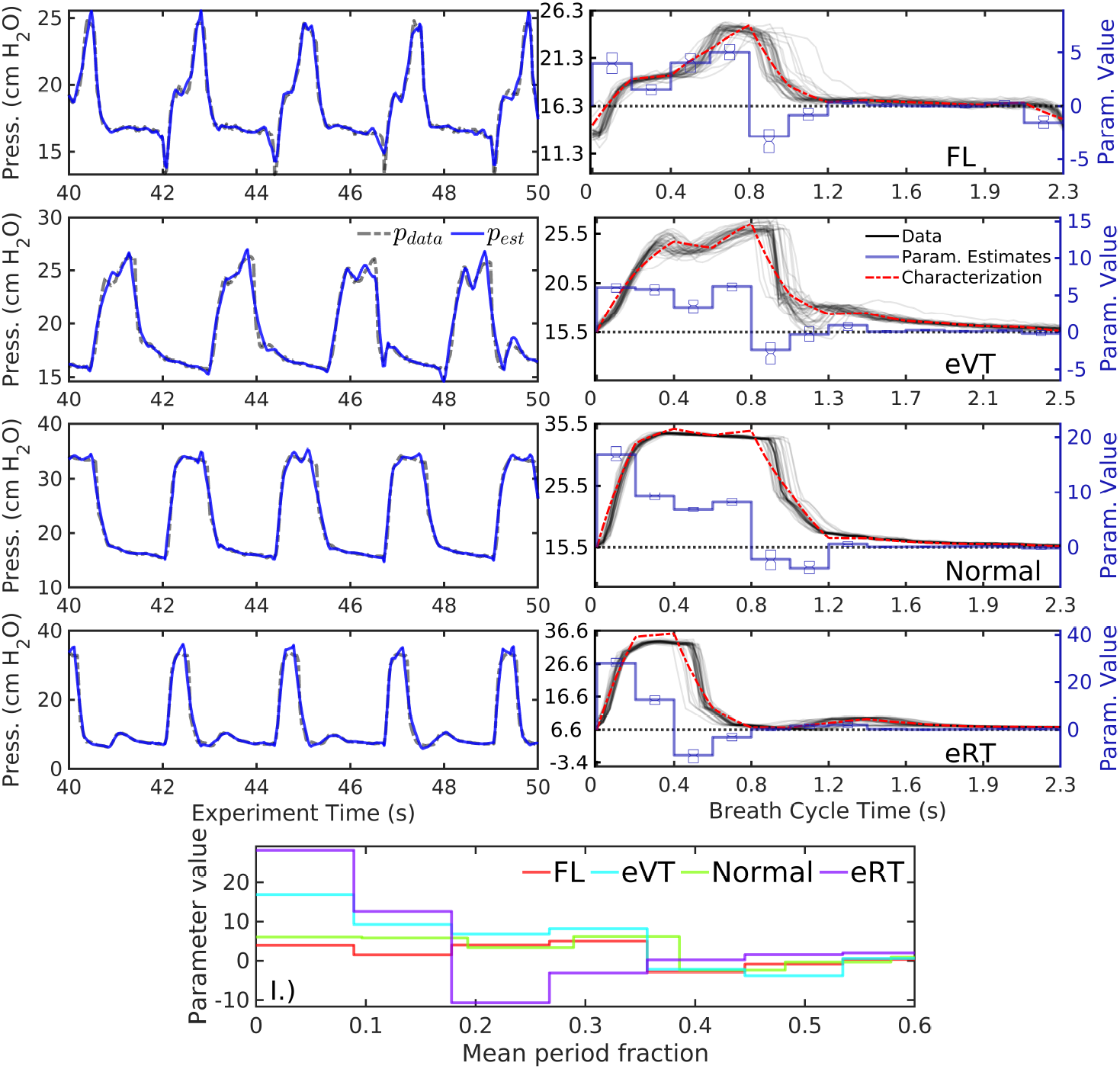
Characterizing patient #2 pressure data with the 12-parameter model *ϕ*_12_, as in Fig3. Differences in the data waveforms have direct, interpretable relationships with differences in descriptive mean parameter sets, which could be used in resolution-dependent LVS breath classification. Summary parameter vectors (I.) are the means of significantly different distributions (*p* < 0.05) and thus distinguishable.

The respective characterizations and parameter averages encode the key differences in waveforms observable in the data and they may be interpreted in parallel ways. For example, inspiration onset pressures (*a*_1_) in normal and eRT sequences (third and fourth rows) are greater than those of FL and eVT sequences (first and second rows). Similarly, the eRT sequence requires a strong negative parameter (*a*_3_) to complete its rapid exhalation. The eVT sequence pressure plateau notch is identified with a decrease in *a*_3_ relative to its neighbors. Late-breath pressure rises during expiration also are evident in eVT and eRT breaths, and these are captured in transient positive values of *a*_7_.

Estimated *pV* loop structures for patient#2 are presented in Fig5) for various choices of *M* governing the parameter resolution. Among the *pV* model resolution tested, the 24-parameter model (*ϕ*_24_) provides the best overall *pV* fit as measured by RMSE over all four sequences. However, *ϕ*_30_-based characterization more accurately represents eVT (B) and normal (C) sequences. The static eRT trace (D) has inaccurate plateau pressure and expiratory volume at this same resolution

**Figure 5:**
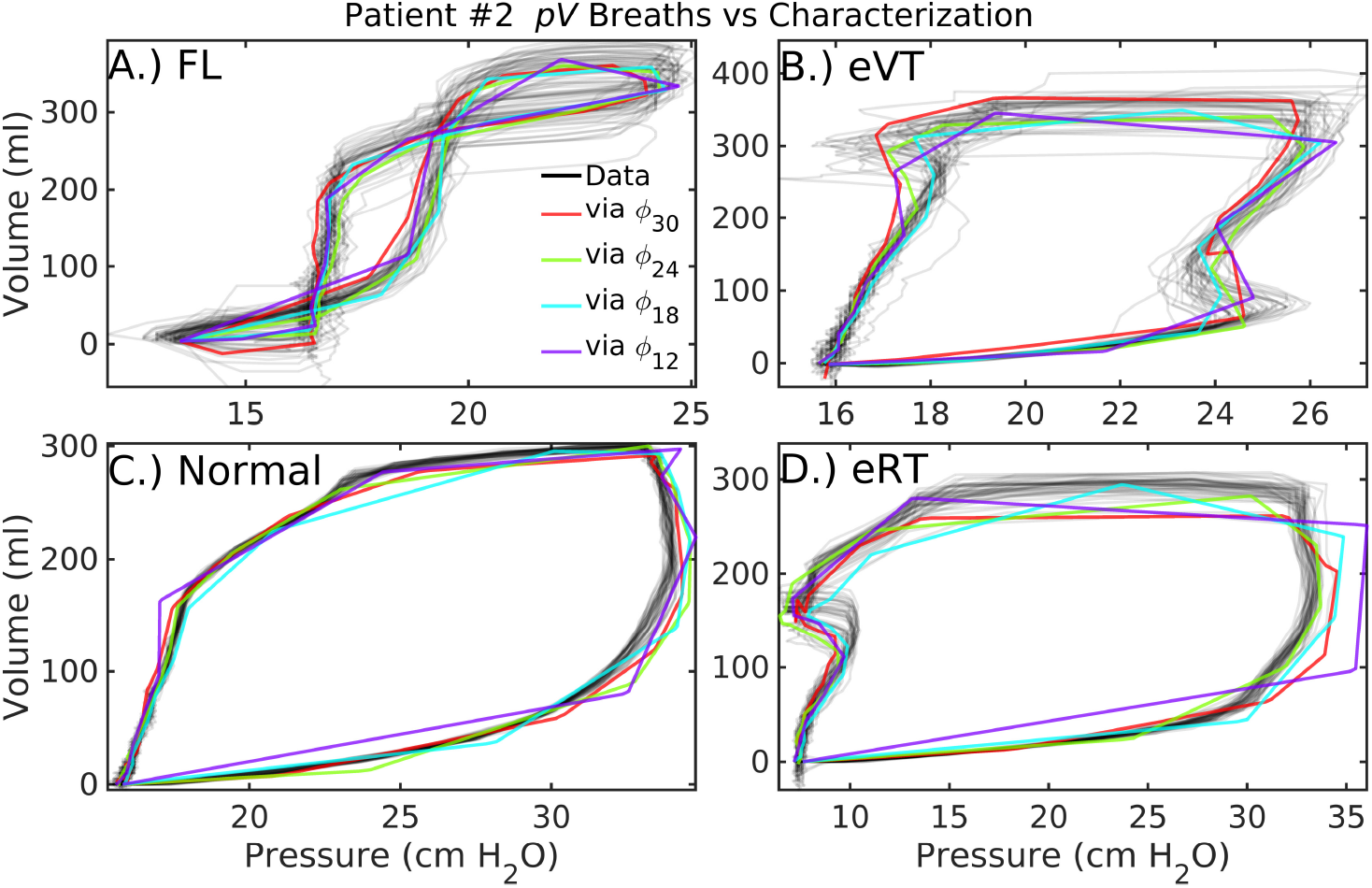
Patient#2 *pV* loop characterizations at various parameter resolutions. In addition to normal breaths (C) similar to those of patient#1, these patient data feature flow-limited breaths (A), early ventilator-termination breaths (B), and early reverse-trigger breaths (D). Increasing model resolution generally improves parametric characterization of the *pV* data, although accuracy also depends on the timing of features in the data and summary window length,

(Table 1). Optimal resolutions notably differ between pressure and volume variables. Further, the eVT pressure sequence for patient#2 (Fig4, D) is adequately represented with 12 parameters to distinguish it from the normal type (F) for phenotyping. Meanwhile, the corresponding *pV* estimate at that resolution (Fig5, B in purple) misrepresents the net compliance, as approximated in terms of the *pV* slope over inspiration. These results suggest the benefit of different component resolutions when targeting the joint *pV* system, as pressure and volume waveform shapes differ profoundly.

### 3.3. Characterization Fidelity

While inferred parameter distributions provide discrete descriptors of data, the fidelity of the waveform characterizations derived from them is important to evaluate as they are more casually interpretable. Fig6 compares stationary, low-resolution characterizations (*M* = 12) to breath-by-breath estimates of the linear compartment model for the pressure sequences previously discussed. The differences are more pronounced in dyssynchronous breaths sequences (C,D,F) than in normal breaths (A,B,F), with higher accuracy achieved by the characterization. Table 2 presents the errors of these estimates to the continuous-time data at several resolutions, including sequences of breaths from additional patient data.

**Figure 6:**
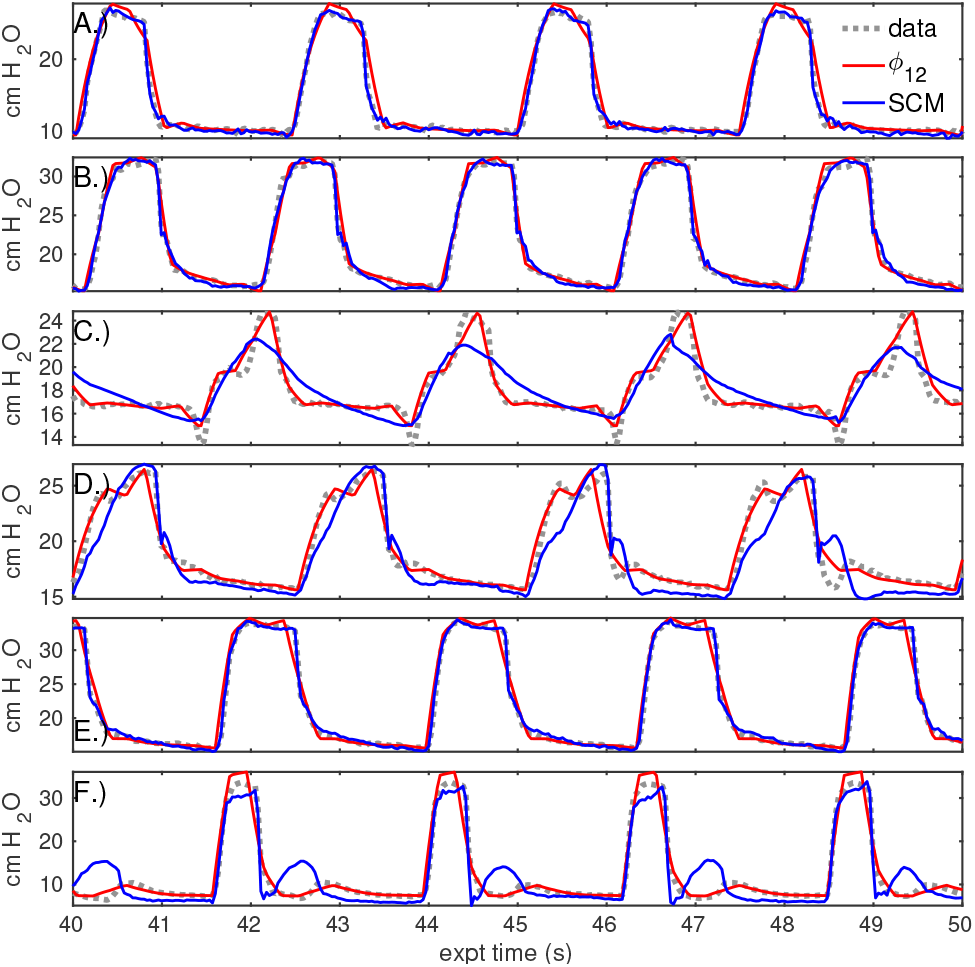
Estimate accuracy of static characterization vs compartment model estimation. The previous set of pressure sequences (grey) are represented by static 12-parameter characterization (red) and by breath-to-breath application of the linear compartment model (blue). Estimation of normal breath pressures (A,B,E) is similar in both models. For breaths with marked dyssynchrony (C,D,F), static characterization outperforms the compartment model. Note that the compartment model estimate is dynamic while the red characterization is stationary; the time axes are regularized to a fixed breath cycle length to plot these signals together.

**Table 2:** Total estimate for various resolution characterizations and the single-compartment model (Eqn(1)), with the lowest value in each row in bold. The large compartment model error for the second sequence of patient #2 arises from a small, persistent difference in phase.

| MODEL | $\phi_{12}$ | $\phi_{18}$ | $\phi_{24}$ | $\phi_{30}$ | Eqn(1) |
| --- | --- | --- | --- | --- | --- |
| Pressure (cm H<sub>2</sub>O) |  |  |  |  |  |
| p#1 Normal 1 | 1.25 | 0.87 | 0.80 | **0.78** | 1.59 |
| p#1 Normal 2 | 0.98 | 0.84 | 0.89 | **0.78** | 1.91 |
| p#2 FL | 0.87 | 0.86 | 0.89 | **0.74** | 3.17 |
| p#2 eVT | 0.93 | 0.82 | **0.74** | 0.79 | 2.23 |
| p#2 Normal | 1.13 | 1.09 | 0.91 | **0.83** | 2.90 |
| p#2 eRT | 1.97 | 2.00 | 1.66 | **1.55** | 4.70 |
| Volume (ml) |  |  |  |  |  |
| p#1 Normal 1 | 8.97 | **8.33** | 8.53 | 9.18 | 20.03 |
| p#1 Normal 2 | 7.75 | **7.31** | 7.62 | 11.56 | 23.23 |
| p#2 FL | 22.05 | **21.73** | 21.78 | 23.36 | 329.23 |
| p#2 eVT | 19.02 | **18.64** | 19.22 | 24.30 | 41.67 |
| p#2 Normal | 10.52 | 8.63 | **8.58** | 10.25 | 33.54 |
| p#2 eRT | 22.45 | 20.98 | **22.33** | 22.44 | 45.09 |

In normal pressure sequences, low-dimensional characterization (*M* = 12) accuracy is comparable to the compartment model and estimates appear qualitatively similar (Figure 6). The relative difference in normal breath errors between the models is typically less than 20%, with several breaths contributing strongly to the compartment error. In contrast, compartmental pressure errors that are 60–70% larger than those of the *ϕ*_12_-characterization for dyssynchronous breaths (panels C,D,F). Sequences of several other patients are estimated SIAppendix D with similar implications.

### 3.4. Summary

The experiment results indicate successful parametric representations of LVS data using informed-model data assimilation. The *O*(10^3^) time-dependent *pV* data points per sequence are reduced to static (2*M* + 2)-parameter descriptions (including PEEP and period) and may be estimated with higher accuracy by either increasing the number of parameters *M* or by shorter summary window lengths. In all cases, inferred parameters accurately represent breath data in continuous time with localized errors related to mis-timings between waveform features and parameter definitions. The patient breaths are distinguishable from the distribution of estimated parameters, even for the generic forcing function used here. These parameters are also interpretable through a priori definition of the forcing function, either as parametric data descriptors or as the waveforms characterizations generated by the forward model. Together, these points indicate that parameter distributions provide a meaningful way to estimate similarity of breaths or breath sequences.

While parameter and characterization fidelity are higher than the compartment model, several factors influence their precision. Parameter uncertainties are greatest where parameter epochs mis-align with waveform features or when waveform feature timings (including the cycle length) vary. The resolution-dependent relationship between discrete epochs and continuous data features influences uncertainty; these factors should be considered when choosing the number of parameters (*M*) and summary window length for an intended application. Increasing the number of parameters may achieve higher characterization fidelity, although this increases model sensitivity to noise and increases the strength of parameter covariance. Further, the statistical summary windows needed to extract characterizations – assumed here to be 40 breaths – may lead to bimodal distributions for which the mean-based characterization is inaccurate.

## 4. Discussion

This work presented a hypothesis-driven approach to modeling the lung-ventilator system (LVS) to infer interpretable parametric representations of human clinical data. Rather than resolve mechanistic processes, the proposed method emphasized model inversion to overcome limitations of the compartment model framework (Sec. 1.1) by combining data- and model-driven methods. To increase key inference attributes of resolution, inferability, and translatability, it replaced the physiology-based model framework with one able to simulate heterogeneous LVS waveforms. This model is a simple differential equation forced by a user-defined parametric function designed to capture waveform properties in the data-optimized solution. Parameters of this model framework are interpretable through their relationship to forcing function they define and the waveform properties to which they are optimized. In the conducted experiments employing a simple non-specific function, these parameters relate to amplitude changes during intra-breath epochs and represent waveform changes during various sub-phases of inspiration and expiration. Data-informed estimates could be distinguished and interpreted by parameter vectors as well as waveform characterization, the model image of the parameter distribution center.

The inferential model system successfully assimilated both synthetic data as well as sequences of human LVS data, comprising both normal and dyssynchronous breaths from patients with ARDS. Synthetic data experiments (SI Appendix C) indicate dynamic parameter tracking in changes of assimilated data for a variety of waveform shapes, with the some error resulting from parameter resolution and temporal coordination. Human data (patient #1) with nuanced waveform differences showed statistically significant differences in estimated parameter distributions. There was also significant contrast in the parametric representation of more varied waveforms (patient #2), including those reflecting LVS dyssynchrony. These results imply that well-estimated parameters have sufficient resolution and flexibility to group and discriminate sequence of breaths based on distributional similarities. Additionally, parameter averages provides adequate, distinguishable representations of data under the assumption of stationary breath cycle length, and are easily extended to characterizations of both waveforms and *pV* loop structure. Parametric summaries, or the forward-model characterizations they induce (via *ϕ_M_* ⟼ *x*), provide an effective basis for discretely organizing, comparing, and analyzing waveform data. Coarse estimation with 12 parameters showed error reductions of more than 50% compared with the linear compartment model in dyssynchronous sequences. Examples of estimated data do not exhaust the possible types and severities of LVS dyssynchrony, although they illustrate the flexibility and robustness of the inferential-model approach.

### 4.1. Hypothesis-driven inference

In broad view, this model strategy is an example of hypothesis-driven inferential modeling in which prior knowledge informs the parameter design of the model, and those parameter values are informed by patient data. This hybrid approach is neither purely empirical like machine learning, nor representative of physical or mechanistic processes as in data assimilation. Parameter estimates, therefore, occupy a grey area between those of more familiar methods. Interpretation – of parameter values or waveform characterization – is based on a priori definitions, while the model itself bears no link to the physical data-generating process. Unlike machine learning, however, parameter values and their patient-level differences may be interpreted in relation to the source data and these definitions. Background knowledge may be used to inform parameter definitions, which provide structure to parameter values without influencing them akin to an uninformative Bayesian prior distribution. A large covariance factor (*γ*) in the inference process ensures that parameter values are strongly informed by the data while parameter definitions control the model output space. From this perspective, the prior knowledge defining the parameters acts as a constraint framework for posing testable hypotheses of the model. The method is also generalizable, although this work focused on ventilator-supported respiration. Namely, the parametrization of clinical waveforms is translatable to other domains (e.g., capnography [41] or intracranial hemodynamics [42, 43]) where underlying processes evade efficient or flexible models needed for analysis at longer timescales.

### 4.2. Pragmatic limitations and trade-offs

While parameter and characterization fidelities are higher than the compartment model, important limitations exist on both interpretability and representation errors. Interpretation depends on the definition of the forcing function, which may be tailored to incorporate domain-specific considerations or knowledge. The generic forcing function used in experiments is highly flexibility but generic. Therefore, its parameters could only be interpreted in relation to breath behavior during a certain interval of the breath. More sophisticated parameters, such as those defining specific waveform features or their timings, provide more granular interpretation but may be difficult to estimate quickly and accurately. For example, if feature timings (e.g. the duration of flow phases) are estimable parameters, the time-ordered nature strongly correlates their values and confounds estimation: a change in one value requires compensatory change in others for a fixed breath cycle length. The trade-off between flexibility and interpretability results from the generic forcing function adopted in experiments. To incorporate background knowledge for LVS pressure estimation, parameter design may be refined to target the inspiration phase at higher resolution than expiration based on the inspiration-to-expiration (I:E) ratio.

The statistical summary process over a window of breaths imposes stationarity assumptions with consequences for interpretability and use of estimates to characterize or differentiate breaths. The minimum resolution needed to capture breath-level features is identified by the Nyquist sampling theorem [44]: a number of parameters *M* > 2*θ/τ* is needed to resolve features with timescale *τ*. Increasing parameter resolution improves the overall fidelity of waveform characterization, up to the threshold of fitting observation noise. However, well-estimated parameters may be non-gaussian, generating characterizations with inaccurate features due to variability of breath duration or feature timing. In particular, the averaging process may eliminate features of interest (e.g. patients #5 and #6 in SIAppendix D) that are captured in the sampled parameter estimates. Characterization fidelity increases by shortening the summary window length from 40 breaths (70– 130 secs) to a few (10–15 sec), although this depends on the data. The ideal number of parameters and the summary length thus present bias-variance trade-offs, with application-specific resolution requirements considered in tandem with the desired summary precision, stability of parameter estimates, and temporal granularity.

Tuning the quantity of generic-model parameters (*M*) and stationarity assumptions may not be suitable for some forms of LVS dyssynchrony. Multimodality in estimated parameters may identify specific forms such as reverse-triggered breaths, which may have a particular repeating sequence of different breaths (e.g., a *(normal, normal, reverse trigger)* breath cycle). In this case, time-ordered parameter estimates are required to discern the sequence from one with a complete transition from normal breaths to reverse-triggered ones. Robust statistical description, such as non-parametric discrimination analysis [45], may be necessary to accurately summarize non-stationary breath sequences from multivariate parameter estimates with correlated components.

### 4.3. Informatics-minded applications

The purpose of encoding LVS waveforms into discrete parameters is for use in research informatics applications involving machine learning and other automated methods. The inference system presented defines vectors to serve in learning tasks such as identification and classification in conjunction with domain-specific information. For studies focusing on individuals, it permits the algorithmic analysis of breath progression in relation to adjunct patient record information not accounted for in parameters. Patient status, intervention, and lung response compose a complete description of respiration within the health care processes, a representation needed to separate lung changes from commingled signals.

Changes in breath behavior occurring distinctly from concurrent changes in ventilator settings and sedation may indicate the effects of ventilation on the lungs. Similarly, parameter trend analysis over hour timescales may differentiate VILI associated with the duration of ventilation from injuries arising from patient efforts such as coughing or voluntary breaths. Such studies revealing the sources of VILI and quantifying their contributions may improve the understanding of lung-ventilator interaction and their consequences. Wider, cohort-oriented works in this vein may new pathways to improved lung-protective ventilator management and patient care strategies. However, applications involving inter-patient comparison require normalization of LVS parameter vectors augmented by patient record information. Additional work is needed to identify suitable normalizations and distance functions required for algorithmic comparisons of data summaries comprising parameterized waveforms and peripheral information.

### 4.4. Concluding Remarks

Changes in lung compliance, capacity, and alveolar recruitment affect the time distribution of lung volume and pressure in response to ventilator forcing. Model parameters quantify observed properties of this distribution with definitions detached from process resolution. This fact limits identifying the source of LVS changes, which may originate from material changes in and of the lung as well as changes in ventilator settings and patient state. Practical applications, therefore, require analyzing *pV* characterization within the full patient-data environment, including knowledge of ventilator settings which the experiments here did not consider. Investigation of the relationship between lung physiology and estimated parameters can proceed in tandem with LVS waveform research, such as those of ventilator control mechanisms e.g., [46], as well as the process-resolving mechanistic models. Although such considerations are beyond the scope of this discussion, the hypothesis-informed parameter descriptions can augment those analyses as low-dimensional representations of waveform data by providing both flexibility and interpretability in a discrete form.

## Acknowledgment

JNS thanks CU Anschutz colleagues Y.Wang and M.Şirlanci for thoughtful and helpful clarifications. This work was supported by R01HL151630 (BJS, PDS, DJA).

## Appendix A. Windowed smoothing via ensemble Kalman transform

In linear ensemble Kalman filtering, the correction of the forecast state is defined as a linear combination of *N* ensemble perturbations about the forecast state. In particular, the Kalman update is 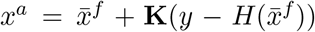 with superscripts *a, f* denoting the analysis and forecast, respectively. The gain matrix **K** may be computed in several different ways, but the full-rank ensemble case always reduces to manufacturing a matrix of the form 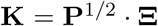, where **P**^1/2^ is a matrix of scaled ensemble state anomalies about the ensemble mean. Importantly, the column space of **K** is determined by the column space of **P**^1/2^. Therefore, the analysis has the form 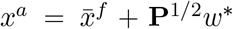 where *w*^*^ is an optimal weight vector for columns of **P**^1/2^ determined by applying the remaining Kalman gain factors 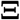 to the data. This implementation is referred to as the ensemble transform as each state *x* in the ensemble span (i.e., the possible EnKF analyses) is uniquely identified with an ensemble expansion coefficient *w* ∈ ℝ*^N^* where *N* is the ensemble size. The ensemble of forecast states is used in the ensemble Kalman filter to approximate dynamical uncertainties. Namely, the ensemble state anomaly matrix **P**^1/2^ is formed column-wise by deviations about their mean and scaled by 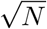; it is a Cholesky factor of an empirical rank-*N* approximation to the true model error covariance **P**. Because error distributions of the model and data are assumed to be gaussian, the analysis *x^a^* may equivalently be identified by minimizing the common cost function

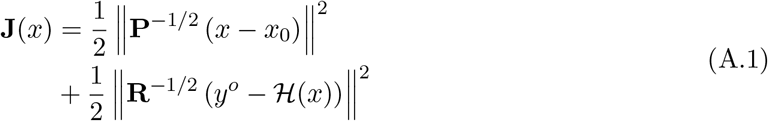

over the *N*-dimensional subspace span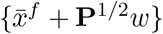. Here, **R**= **R**^1/2^**R**^*T*/2^ is the error covariance matrix associated with the observation process, which includes error and uncertainties involved in translating model states to equivalents of observational data as well as instrument errors when generating observational data from the real world. The inverse of the hessian matrix associated with **J**(*x^a^*) identifies the posterior error covariance matrix [47]. From a Bayes’ Rule perspective, this optimum is the mode of the posterior probability distribution produced when the forecast model is updated based on new data [48].

## Appendix A.1. Asynchronous Smoothing

Extensions of the EnKF method described above to assimilate observations occurring at various times within a moving window are developed in e.g. [49, 50]. The passage below follows the variational perspective of [51, 52] for conciseness.

The equivalent of Eqn. (A.1) conditioning the optima transform variable *w* on multiple observations 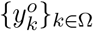 in the window is given by given by the quadratic function:

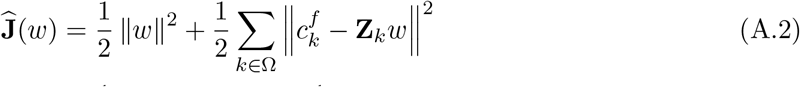

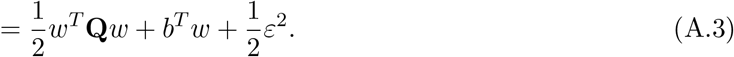

Here, *k* ∈ Ω ⊂ {1*,…, L*} indexes the times associated with observations being assimilated, at which times one requires observed model forecast anomalies **Z**_*k*_ and scaled misfits to the data 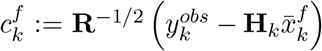. In Eqn.A.3, the object 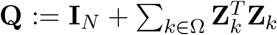 is the hessian matrix for the system linearized about the initial trajectory 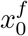, the vector 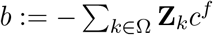 is the linear coefficient, and 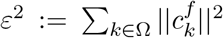 is initial trajectory error. In applications with overlapping-windowed analysis, Ω regulates whether data assimilated in a previous window are re-assimilated in the present one.

A linear approximation to the variational optimum is found algebraically by targeting the mode of the posterior distribution. Specifically, solving 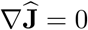 yields the (approximately) optimal coefficient

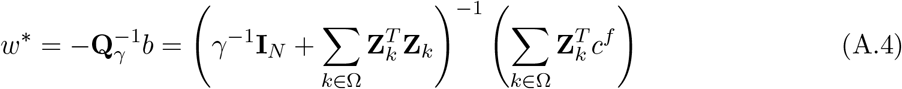

where *γ* is an optional covariance inflation factor with *γ* > 1 increasing relative weight on errors to the data. The true minimum of the nonlinear problem may be obtained by minimizing Eqn.(A.2) by iterating over re-forecasts; Ref. [52] includes a more comprehensive analysis of different schemes for computing the full optimum. Such approaches are important in strongly nonlinear systems whose ensemble of solutions diverges strongly within the assimilation window, which is not the case with the simple model considered in this work.

Once the optimal analysis state *x_S_* is calculated, the ensemble variations about this new distributional center are found by re-weighting the forecast covariance Cholesky factors by those of the quadratic program inverse Hessian matrix: 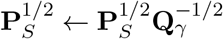.

## Appendix B. Algorithm Parameters for Experiments

**Table B.3:** Non-forcing parameters and experiment details are given, where ‘*data*’ indicates values extracted from the data.

|  | Symbol | Synthetic Expt. | Human Expt. |
| --- | --- | --- | --- |
| Ensemble Size | $N$ | 32 | 60 |
| Sample Rate [Hz] | - | 25 | 32.25 |
| Window Length | $L$ | 75 | 50 |
| Update Step | $S$ | 25 | 20 |
| ODE Parameter | $g$ | $e$ | $e$ |
| Reference State | $x_0$ | 20 | data |
| Breath Cycle | $\theta$ | 5 s | data |
| Initial Parameters | - | 0.5 | 0.5 |
| Data RMSE | $\mathbf{R}^{-1/2}$ | 5% | 5% |
| Parameter RMSE | $\mathbf{P}_0^{1/2}$ | 0.025 | 0.025 |
| Covariance Inflation | $\gamma$ | 100 | 100 |

## Appendix C. Parameter accuracy via synthetic data experiments

Parameters were changed every 8 breaths in the 10-parameter model (*ϕ*_10_) with a fixed 5-second cycle length to generate two minutes of 25 Hz synthetic data (FigC.7, top panel in grey). Inference experiment results (blue, throughout figure) use a high covariance inflation factor (*γ* is *O*(10^2^)) to sharply focus the solution on a noise-contaminated version of these data. Simulated waveforms accurately reconstruct window-wise trajectories (top, blue) from local parameter estimates inferred from noisy data. Estimated parameter values (lower, blue) accurately and quickly track changes of the true parameters defining the data (lower, grey), with transitions to new parameter values occurring within two update windows (twice the length of *t_S_*, or about 2 seconds).

Early breath parameters change sooner than parameters later in the breath, seen by comparing transitions of *a*_1_ and *a*_2_ to those of *a*_7_ and *a*_8_ (around 60 seconds). This is a result of early-breath data begin assimilated sooner, with upstream parameter dependence also evident in the estimates. For example, the value of *a*_1_ is slightly underestimated for *t* ∈ (30, 60) and the value of *a*_2_ is correspondingly slightly overestimated to compensate. Relative errors are smallest for large magnitude parameters and largest for those with values near zero (such as *a*_10_). The large error in estimated *a*_8_ for *t* ∈ (60, 90), where the true value of 1.8 is estimated as ≈ 2.8, is a consequence of diminished model sensitivity to smaller-magnitude parameters. The inversion process accurately infers the parameters dominant in determining the solution shape, aptly tracks parameter set changes in time, and thereby gives dynamic representations of the data via model parameters.

## Appendix D. Estimation of Additional Patients

This addenda estimates several breath sequences in addition to those of Sec3. The estimated sequences from three patients (#3–#6) were selected for a minimum of data artifacts and consistency of waveform behavior. In these sequences, the windowed model inversion tightly fits the parametric model to data (FigD.8, left column) regardless of the waveform shape. Characterizations (right, red) give low-dimensional representations including resolution-dependent inaccuracies. For example, the pressure peak of patient#3 sequence 2 (second row) is highly variable in both amplitude and timing; the associated characterization based on the mean is inaccurate, but the variability is captured and encoded into the parameter uncertainty (left, blue). Similar inaccuracy is evident in the late pressure plateau pressure peak of patient #5’s sequence (fourth row). Rather than increasing the number of parameters, resolving it requires decreasing the summary window to 4–5 breath windows. The pressure waveforms of patient #6’s sequence (bottom row) have an irregular cycle, leading to a bimodal parameter estimate when assuming a common period for the 40 breaths. This generates large uncertainties in all estimated parameters, and the sequence is best characterized by the coarse resolution model. Nevertheless, static characterizations based on the mean parameter vector generally have lower RMSE over the 40-breath sequence than corresponding estimates made by the linear compartment model (TableD.4). The first sequence of patients is better estimated by the compartment model than the low-resolution characterization (*M* = 12); for all other sequences, errors decrease by over 40% using the 12 parameter model (FigD.9).

**Figure C.7:**
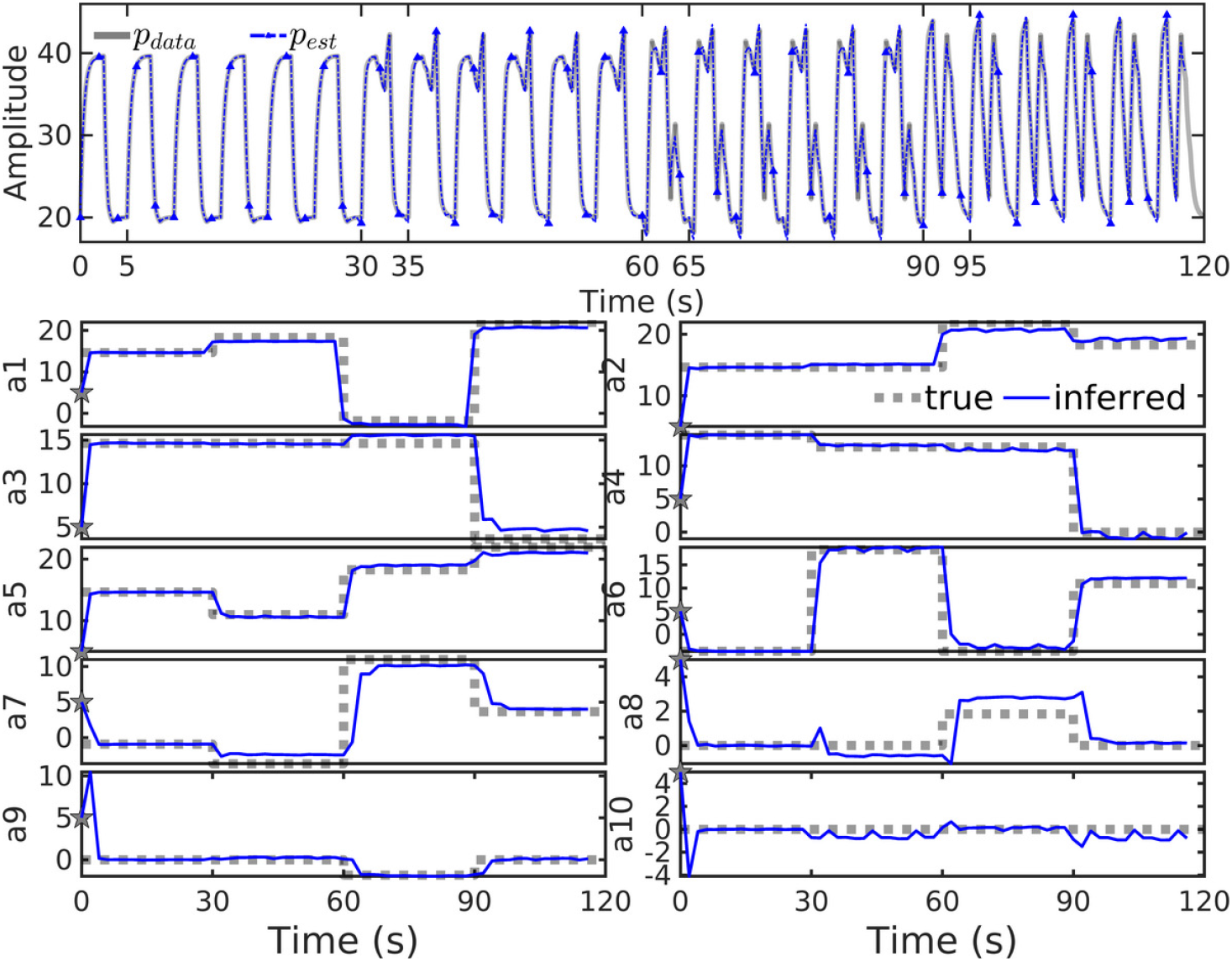
Inversion of *ϕ*_10_ with synthetic data (upper panel, grey) generated by known parameters (lower panels, dashed grey) correctly identifies the correct parameter values and their changes in time. Reconstructed waveforms (upper, blue) accurately track the correct solutions.

**Figure D.8:**
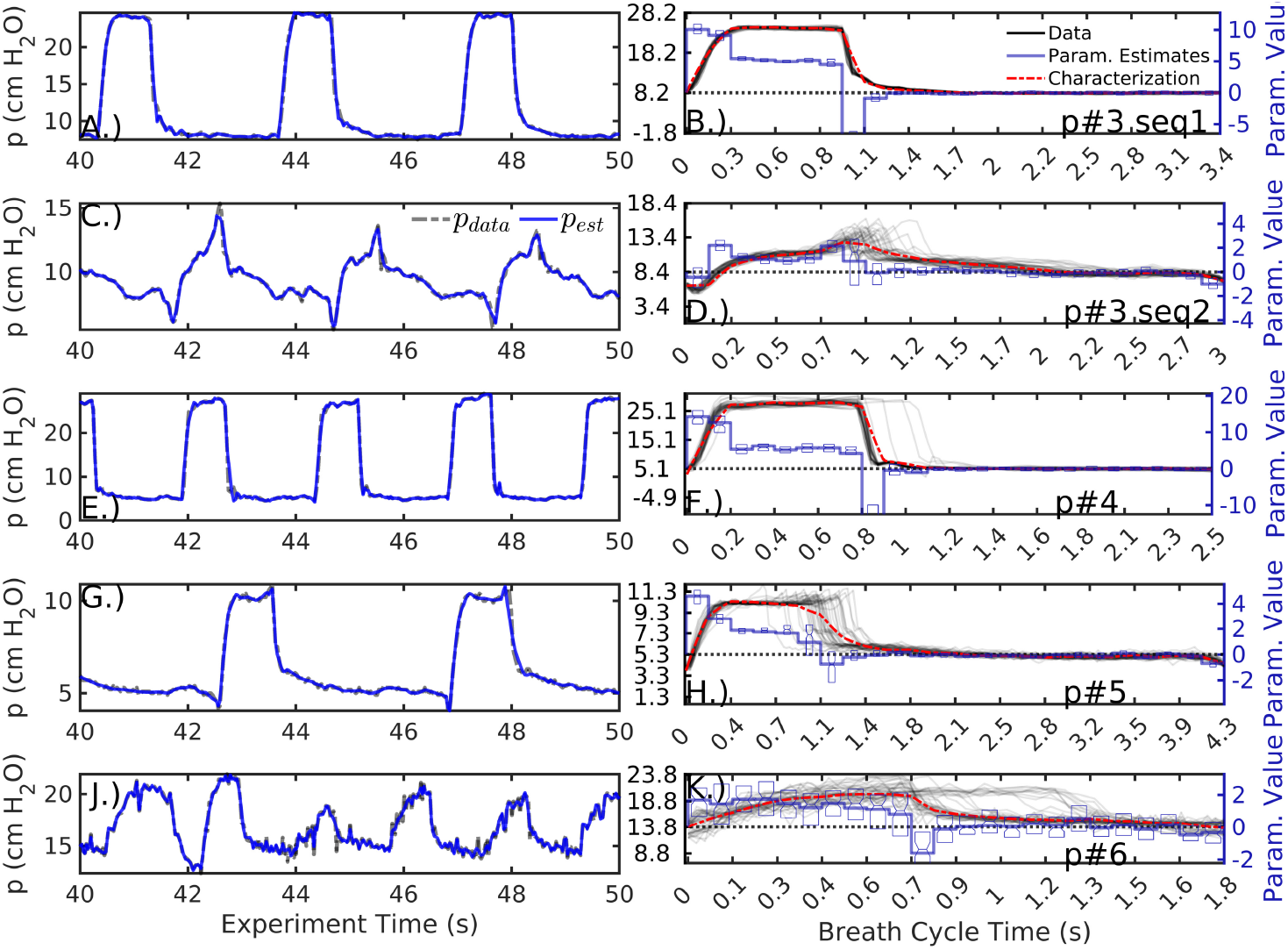
Parameter estimation and characterization (*M* = 24) of additional pressure sequences, as in Figs3,4. Specific dyssynchrony types are not identified but breaths in the first and third row are relatively normal. The last row features breaths with an irregular cycle; the data plotted in panel K indicate a bimodal breath cycle.

**Table D.4:** RMS errors (RMSE) of estimates to data for various resolutions and the single compartment model (Eqn(1)), with lowest error among proposed static model characterizations is indicated in bold. Large errors in volume of compartment model estimation result from persistent differences in phase.

| MODEL | $\phi_{12}$ | $\phi_{18}$ | $\phi_{24}$ | $\phi_{30}$ | Eqn(1) |
| --- | --- | --- | --- | --- | --- |
| Pressure (cm H<sub>2</sub>O) |  |  |  |  |  |
| p#3 seq1 | 1.03 | 0.80 | **0.60** | 1.11 | 0.65 |
| p#3 seq2 | 0.70 | 0.66 | 0.63 | **0.62** | 1.43 |
| p#4 | 2.61 | 2.27 | 2.06 | **1.87** | 4.71 |
| p#5 | 0.63 | **0.62** | 0.63 | 0.63 | 3.32 |
| p#6 | **1.50** | 1.51 | 1.57 | 1.97 | 3.46 |
| Volume (ml) |  |  |  |  |  |
| p#3 seq1 | 16.92 | 12.11 | 10.07 | **9.70** | 23.81 |
| p#3 seq2 | 56.55 | 55.83 | 55.80 | **55.60** | 349.77 |
| p#4 | 29.31 | **29.09** | 29.59 | 31.37 | 58.43 |
| p#5 | 44.02 | 43.76 | 43.71 | **43.70** | 230.45 |
| p#6 | **148.32** | 148.73 | 155.00 | 166.66 | 262.49 |

**Figure D.9:**
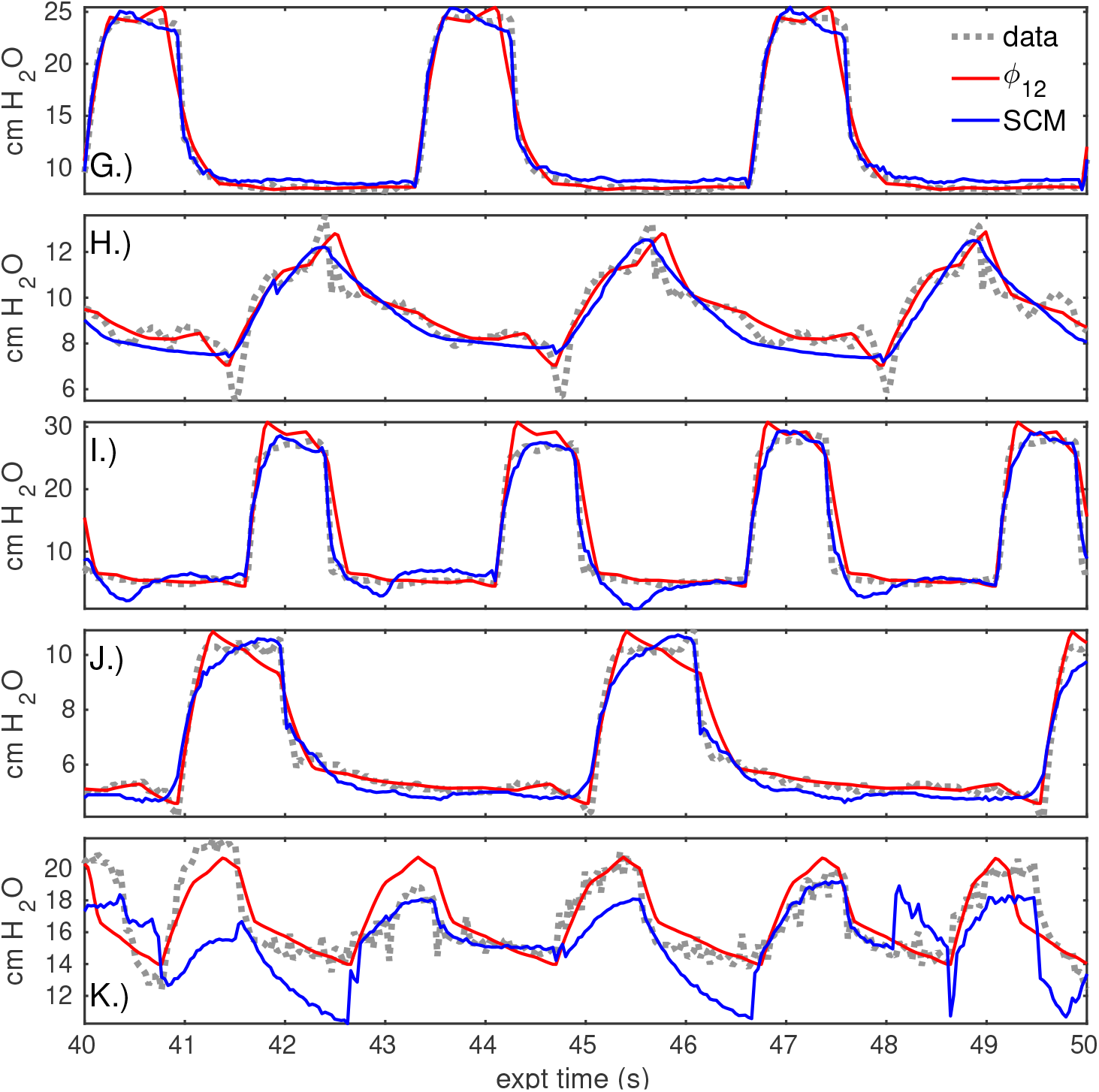
Compartment model estimate and static characterization (*M* = 12) compared to pressure data, as in Fig6. The rows correspond to the rows of the previous figure and of the following table. The time axis of each sequence is regularized to a fixed breath cycle length to depict these signals together.

